# Perturbation-based gene regulatory network inference to unravel oncogenic mechanisms

**DOI:** 10.1101/735514

**Authors:** Daniel Morgan, Matthew Studham, Andreas Tjärnberg, Holger Weishaupt, Fredrik J. Swartling, Torbjörn E. M. Nordling, Erik L.L. Sonnhammer

## Abstract

The gene regulatory network (GRN) of human cells encodes mechanisms to ensure proper functioning. However, if this GRN is dysregulated, the cell may enter into a disease state such as cancer. Understanding the GRN as a system can therefore help identify novel mechanisms underlying disease, which can lead to new therapies. Reliable inference of GRNs is however still a major challenge in systems biology.

To deduce regulatory interactions relevant to cancer, we applied a recent computational inference framework to data from perturbation experiments in squamous carcinoma cell line A431. GRNs were inferred using several methods, and the false discovery rate was controlled by the NestBoot framework. We developed a novel approach to assess the predictiveness of inferred GRNs against validation data, despite the lack of a gold standard. The best GRN was significantly more predictive than the null model, both in crossvalidated benchmarks and for an independent dataset of the same genes under a different perturbation design. It agrees with many known links, in addition to predicting a large number of novel interactions from which a subset was experimentally validated. The inferred GRN captures regulatory interactions central to cancer-relevant processes and thus provides mechanistic insights that are useful for future cancer research.

**Data available at GSE125958:** Inferred GRNs and inference statistics available at https://dcolin.shinyapps.io/CancerGRN/ Software available at https://bitbucket.org/sonnhammergrni/genespider/src/BFECV/

**Author Summary:** Cancer is the second most common cause of death globally, and although cancer treatments have improved in recent years, we need to understand how regulatory mechanisms are altered in cancer to combat the disease efficiently. By applying gene perturbations and inference of gene regulatory networks to 40 genes known or suspected to have a role in cancer due to interactions with the oncogene MYC, we deduce their underlying regulatory interactions. Using a recent computational framework for inference together with a novel method for cross validation, we infer a reliable regulatory model of this system in a completely data driven manner, not reliant on literature or priors. The novel interactions add to the understanding of the progressive oncogenic regulatory process and may provide new targets for therapy.

## Introduction

Cancer can be seen as an altered state of the regulatory systems that control cell proliferation and cell death. Such systems are generally not sensitive to individual gene malfunctions, but an aggregation of aberrations can lead to sufficient dysregulation to cause cancer. Reliable models of these regulatory interactions would offer insight into key mechanistic alterations for therapeutic targeting. Cancer subtype-specific gene regulatory networks (GRN) encode intracellular dynamics [1], and offer understanding into the functional changes driving disease development. Inference of such models generally exploits certain aspects of the experimental setup, such as pooling among replicates to amplify signal, or makes use of prior knowledge [2,3]. The experimental techniques, setup, and data collection quality determine the quality of an inferred GRN model. However, practical limitations of experimentation, such as high noise levels and few experiments relative to the vast combinatorial landscape of possible regulatory interactions, often prevent any GRN inference methods from inferring a correct GRN[4]. Methods using data from known and systematic perturbations have shown greater accuracy among inference techniques since more information is available to determine regulatory causal mechanisms in the system[5].

GRN inference has proven its value to unravel novel regulatory links of biological significance. For instance, ARACNe was applied to gene expression profiles to predict a glioma-specific GRN, revealing that C/EBPbeta and STAT3 are master regulators of mesenchymal transformation, which was validated experimentally [6]. In another study, eight key genes were knocked down by siRNA, and the gene expression together with prior knowledge were used to infer a GRN network in the RAS pathway with good validation performance [7].

A large number of GRN inference algorithms exist. In a survey by the DREAM5 project [8] of 35 inference methods, it was shown on simulated and *E*. *coli* data that some methods performed better than random predictions. However, many methods did not outperform random, and on yeast data no method performed much better than random. Since this study, the community has developed methods for integrating various priors: literature/database, ATAC-seq, DNase I hypersensitive sites, ChIP-Seq, or protein data to increase information about the system [9–11]. The categorization of methods has expanded too, most notably to include machine learning or neural models, adding some 15 more methods to the DREAM5 survey [12]. The benchmarking suite NetBenchmark [13] was developed to provide the community with direction in choosing a best suited method for doing inference. However, this failed in arriving at a universal method which outperformed all other methods across all datasets. Some methods perform generally poorly In benchmarks, for instance ARACNe2 [14] performed only marginally better than random with an average AUROC of 0.58 across different sized networks, which is consistent with another benchmark study [15]. These benchmarks and surveys testify to the fact that network inference remains a very challenging task considering expression data alone. Integrative approaches can improve performance but depend on the availability of different types of omics data, and faces challenges such as varying experimental setups, heterogeneity, and quality of input data. In many cases, only expression data are available however, and here the importance of data quality is paramount In this study, we deployed perturbations through siRNA gene knockdown of each gene in our literature-curated set of cancer-related regulator genes, each followed by transcriptomics measurements of all genes, in order to measure the global influence of each individual gene. Knockdown experiments are more informative about the system than irreversible and complete knockout which may cause drastic rewiring of the network into another system. Assuming a linear time invariant (LTI) system [17], once the system has reached a steady-state, a GRN can be inferred by solving a set of first order ordinary differential equations (ODEs)[18] in the form of our linear model (eq.1). Importantly, however, our linear model is reliant upon a known perturbation design, which additionally informs our model. In this way a selected set of 40 genes relevant to cancer were perturbed, and the transcriptomic response data were used to construct a model of underlying regulatory interactions. We inferred these interactions by relating the effect of the gene perturbations to the expression of the readout genes, using three GRN inference algorithms well suited for employing our linear model and perturbation design: LASSO, LSCO, and TLSCO.

A drawback of all GRN inference algorithms is that they generally produce erroneous GRNs if the noise level is high [15,19]. To ensure inference of reliable GRNs, we employed NestBoot [16], a recent framework implementing nested bootstrapping, wrapped around any individual GRN inference method to better account for sample variation and noise. Contained within the GeneSPIDER package [15], NestBoot generates bootstrap support distributions for links inferred from measured as well as shuffled data[20], and minimizes false links by comparing them. This way Nestboot is able to discard links even if they have high bootstrap support, if they also have this in the null distribution. NestBoot has been shown to give substantially increased inference accuracy across both synthetic and experimental datasets when compared to the methods in their native implementation.

In order to measure the accuracy of an inferred GRN, a true GRN is required. Because we lack a true GRN in the case of real data, we here introduce methodology to assess the predictiveness of an inferred GRN. Note that we are not presenting a GRN inference method on its own but is rather a way to assess the quality of a given GRN. We first use it to measure a GRN’s ability to predict the data compared to a distribution of GRNs with the same topology as the inferred one but whose links have been shuffled. We complemented this performance evaluation by measuring the GRN’s ability to predict the data compared to a distribution of shuffled data. Finally, we present the best performing GRN in detail, although the other inferred GRNs are largely subsets of each other and mostly perform well too. Two of the novel links of the best GRN were experimentally validated. The presented GRN captures regulatory interactions central to cancer-relevant processes and we foresee that it can provide mechanistic insights that can help to guide future cancer research. For instance, many cancers are caused by dysregulation of the MYC oncogene, hence our finding of a new regulator of MYC may potentially lead to new therapies.

## Methods

### GRN inference

The fold change is calculated in comparison to the spike-in for all knockdown experiments. It is used in combination with the collective experimental design matrix (describing the location of perturbed and readout genes) to determine the GRN, i.e. the interaction matrix A, of regulatory effects from gene *j* to *i* in element *a*_*ij*_. We use a linear ODE model, similar to [17,21], which simplifies to

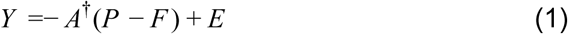

where Y is an expression matrix of calculated fold changes, with N genes (rows) and M experiments (columns). In Eq. 1, *P* is the design matrix if we solve for *A*^†^, where the Moore-Penrose generalized inverse, denoted *†*, is used throughout in place of the inverse due to computational intractability wherein sparse GRNs might be rank deficient. However as we want to solve for *A* and not *A*^†^, we reformulate Eq. 1 to a traditional regression problem on errors-in-variables form, - (*P* - *F*) = *A*(*Y* - *E*). The error in *Y* and *P* are represented as measurement error *E* and process error *F*, respectively, as defined in Table S1. *F* is used as an estimate of the variation in the perturbation, *e*.*g*. siRNA efficiency or environment, while E is used as an estimate of the variation inherent to the cells’ expression as well as error in plate reading [26].

Three methods are employed to perform model selection and parameter estimation simultaneously. LSCO (least squares with a cut off to produce variably sparse networks)[22] was chosen for its resemblance to the standard ordinary least squares method, LASSO (least absolute shrinkage and selection operator)[23] was chosen for its proven ability to find the sparse solution with minimum errors, and TLSCO (total least squares [24] with the same sparsity inducing cut off as LSCO) for its ability to model error in both the dependent and independent variables [19]. Each method is encapsulated within the nested bootstrapping framework to estimate the linear model in an accurate and reproducible manner by limiting false discovery rate (FDR), in their native configuration.

### GRN validation without Gold Standard

To evaluate the goodness-of-fit of the inferred network in a prediction error framework one needs to balance the measurement and process errors. This optimization occurs during the leave one out procedure (Algorithm 1), using the CVX convex optimization package (v1.22)[25] for MATLAB, where the left out gene (g) is expressed as a linear combination of the other experiments (crossvalidation). The aim of this procedure is to balance the measurement and process errors when predicting the left out gene using crossvalidation.

#### Algorithm 1 BalanceFitError (BFE) with Crossvalidation

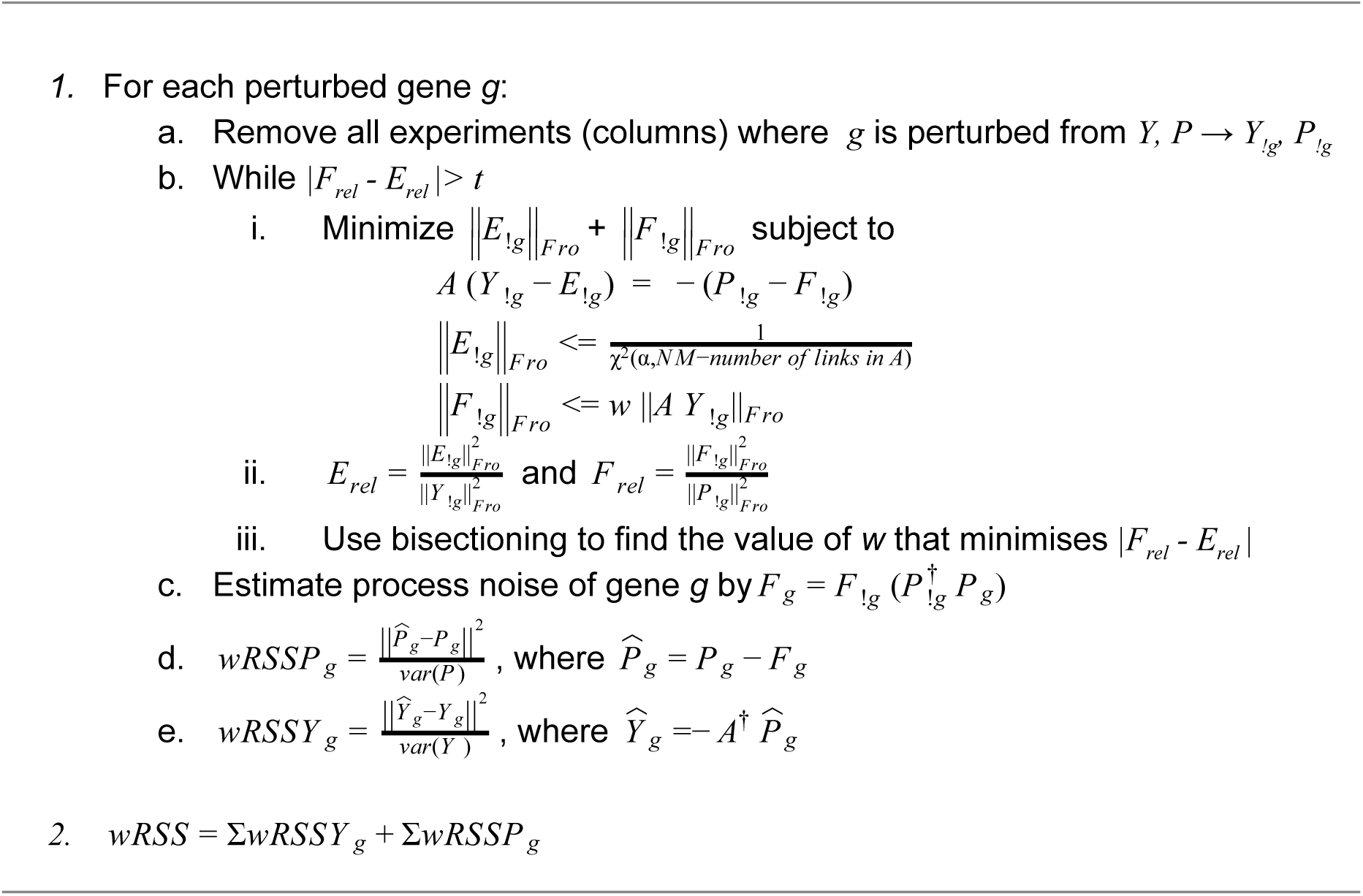

In the BalanceFitError algorithm, which estimates the generalization error - a proxy for the prediction error without knowing the true GRN. *A* contains the inferred GRN structure (topology), with each non-zero value representing an interaction and each zero a lack of interaction, i.e. pseudo-direct influence [17]. For each perturbed gene one estimates the perturbation and response based on the balanced measurement and process errors of all other genes and compare it to the intended perturbation and observed response. The noise of *Y*, represented as *E*, is used as an estimate of the variance inherent to the cells’ expression as well as error in plate reading[26]. Since error is a function of the degrees of freedom of the given matrix, relative error (*E*_*rel*,_ *F*_*rel*_) is used to more equally balance these errors. From step i. to iii. all matrices have the perturbation experiments of the left out gene *g* removed and are thus denoted *!g*. However, *A* maintains all genes, remaining square throughout, and later the left out experiments can be predicted from the remaining data. This method to evaluate the goodness-of-fit is used on all inferred networks. All inference methods used here have a regularization parameter that determines the number of nonzero parameters in the models, which is varied to span the complete range from empty to full network.

Because our leave out procedure assesses individual gene prediction errors, we assembled null GRN performance distributions by shuffling GRN links and fitting these new networks to the data to create both a standardized and fairly conservative link weight using a constrained least squares (CLS) algorithm [22,27]. To this end we implement a Monte Carlo sampling method, sampling links to maintain the node indegree and preserving hubs thereby approximating an estimated link null distribution based on the inferred GRN judged to generate conservative and fair null GRNs. For a fair comparison, both the inferred and shuffled GRNs are fit to the original data. However topology and sign are preserved in the GRNs. To obtain a measure of the goodness of fit of both inferred and shuffled GRNs, crossvalidation was used to calculate the weighted residual sum of squares (wRSS) of the original training data while balancing the measurement and process errors as described in Algorithm 1. We are able to predict a left out gene in step *c* and *d* by expressing it as a linear combination of the other genes. This goodness of fit measure was also made of the inferred GRN’s ability to under crossvalidation predict the original data compared to the distribution of prediction errors using shuffled data. The relative error metric comparing measured and shuffled wRSS (Fig. 2, 3, S3, S4) is complemented by R^2^ values (Fig. S6). Before calculating these, each GRN parameters were modified to ensure that the predicted response remained similarly bounded as the observed gene expression. This was done by performing singular value decomposition of the GRN, setting singular values below a cutoff to zero, and then reconstructing the GRN without the smallest singular values. This GRN was then fit to the training data under crossvalidation to generate predicted expression responses in the same way as described above. The cutoff on the minimum singular value was set independently for each GRN to ensure that the predicted expression values were within the range of the measured values. The small singular values generally represent noise if the data is ill-conditioned and removing them reduces the effect of noise.

**Fig. 1.**
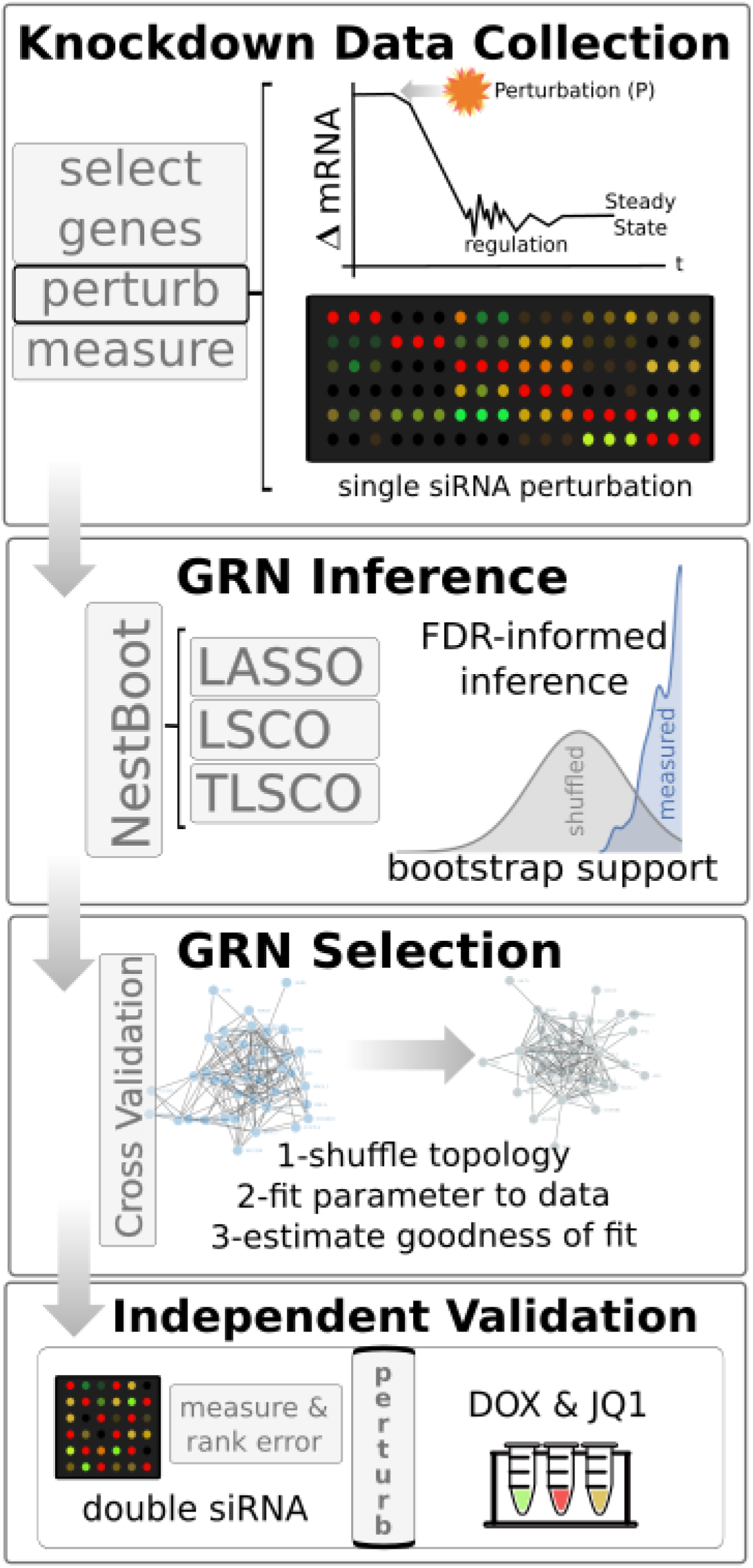
Workflow of project. siRNA perturbation experiments were carried out systematically per gene, resulting in a change in mRNA level which elicits a regulatory response over time before reaching steady state when gene expression was measured. GRN inference: Nested bootstrapping was applied to three inference methods, producing GRNs with an FDR set to 5%. GRN Selection: Estimate goodness of fit under crossvalidation and error balancing, and compare to shuffled topologies (Algorithm 1). Independent Validation: Each inferred GRNs’ ability to predict an independent dataset was evaluated in comparison to a distribution of shuffled topologies. Finally, the overall most predictive GRN was selected. Two novel links were experimentally validated.

**Fig. 2.**
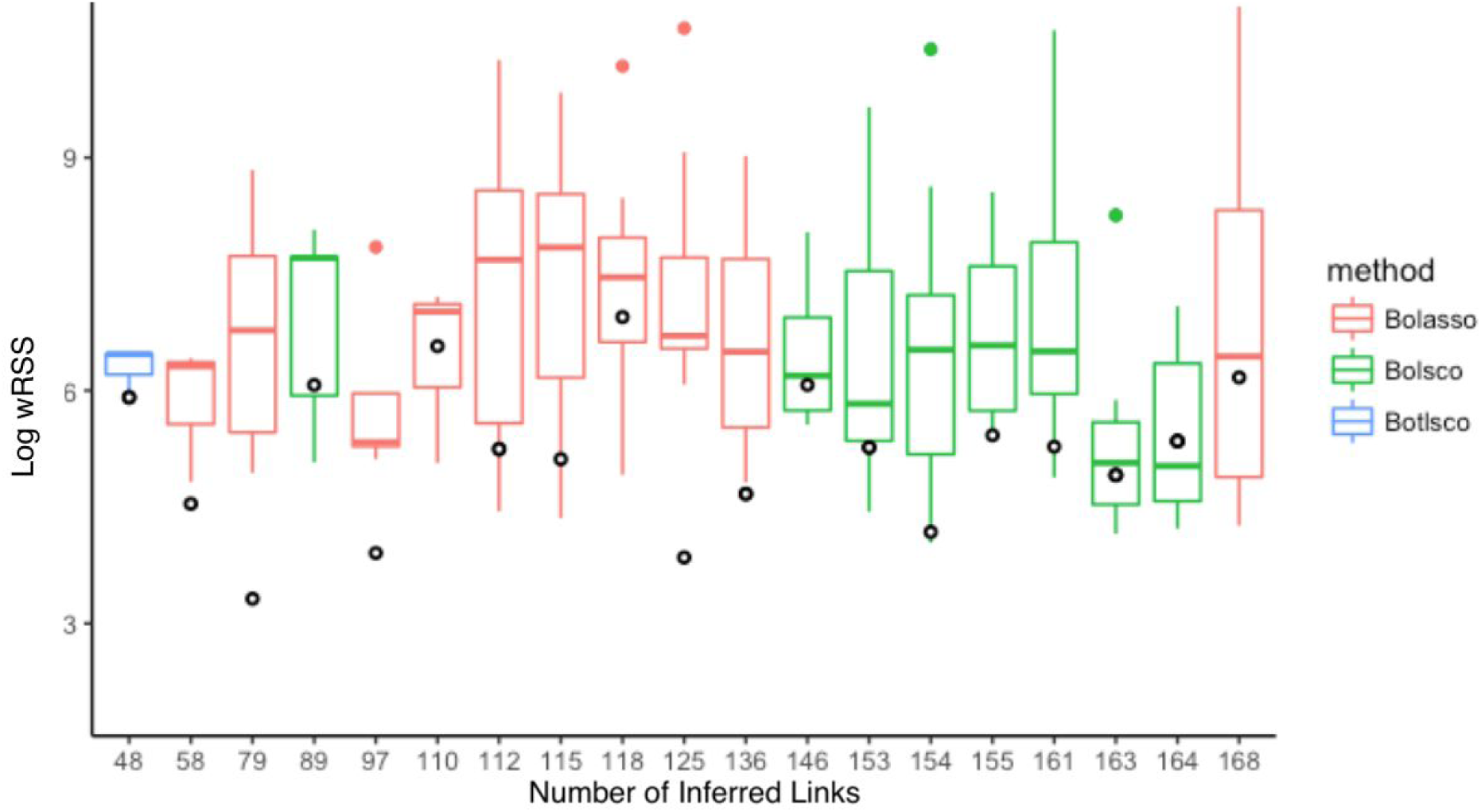
Validation of inferred GRNs’ topologies. Each x-axis tick mark shows the prediction performance in terms of the wRSS error of each inferred GRN topology (circles) fit to training data under crossvalidation, compared to its shuffled topologies. The box displays the median and interquartile range, and whiskers bound points maximally extending 1.5 times this range. Beyond this, outlier points are shown.

**Fig. 3.**
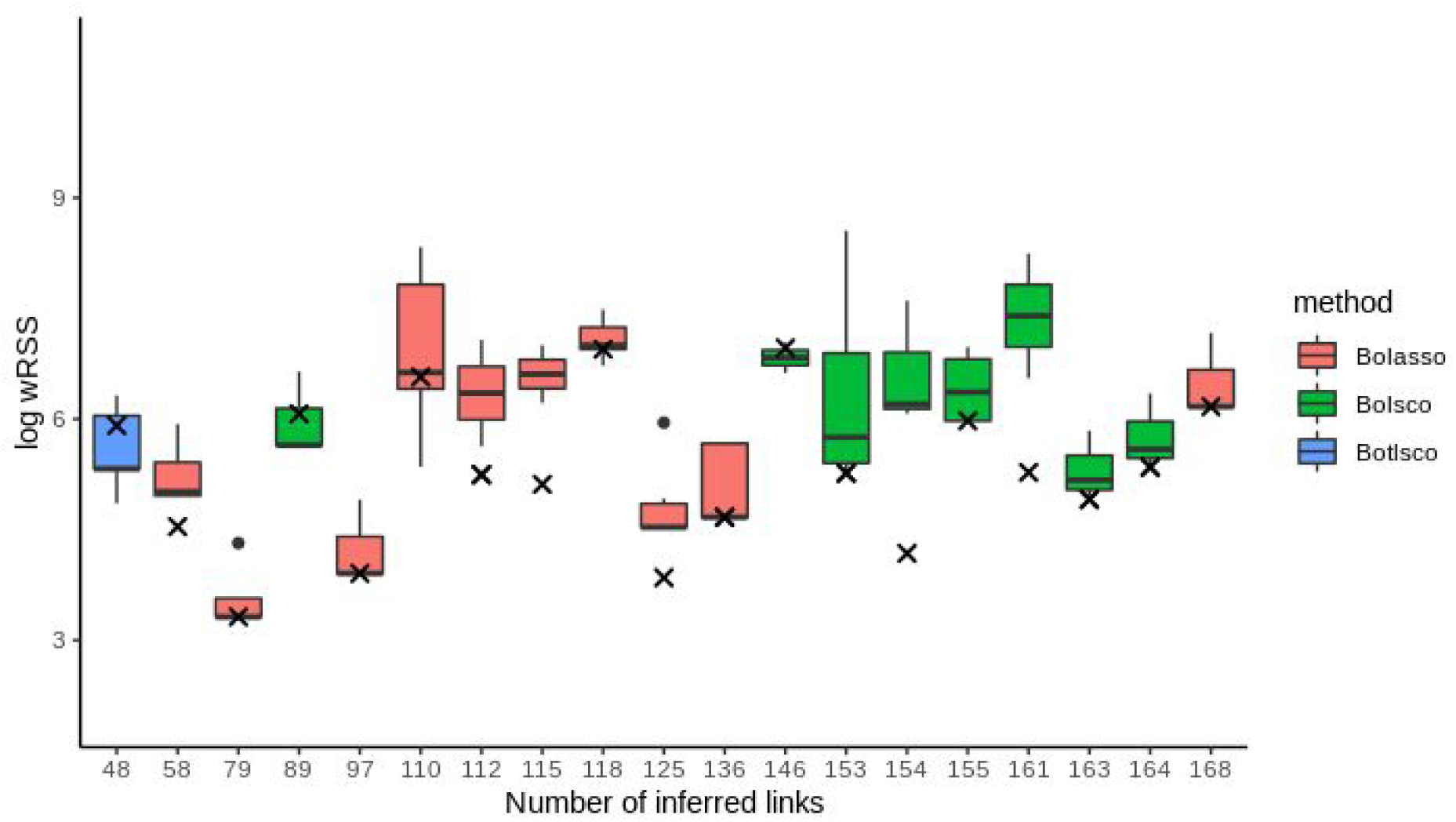
Validation of the inferred GRNs’ fit to the measured data. Each x-axis tick mark shows the prediction performance in terms of the wRSS of an inferred GRN topology fit to training data under crossvalidation, compared to its ability to fit shuffled data. X marks represent the inferred GRNs. The filled color box displays the median and interquartile range, and whiskers bound points maximally extending 1.5 times this range. Beyond this, outlier points are shown.

To further verify both predictiveness and generalizability, these GRNs are also applied to a second independent, validation dataset based on the same genes knocked down in pairs, in single replicates. While this data is not used to infer GRNs, we apply the same crossvalidation strategy as for the original data to validate the GRNs. This is necessary for parameter fitting, since the process error is different from the single knockdown data. Furthermore, by running the same pipeline we obtain a comparable measure of how well the independent data fits our inferred GRNs, and build null distributions of expected error from shuffled GRNs to examine an inferred GRNs’ ability to predict the data.

## Results

A four step procedure was implemented to generate a cancer-centric GRN oriented towards the MYC oncogene (Fig. 1). First, a list of 340 cancer associated genes was ranked heuristically by what was known of each, giving preference to genes with known associations to both cancer and MYC. The criteria were as follows, in decreasing order of rank: i. members of a complex with MYC affecting or not affecting transcripts, ii. genes directly affecting MYC or MYC transcripts (activating/repressing), iii. genes affecting MYC targeted transcripts, and iv. genes indirectly affecting MYC transcripts. Only genes expressed in the used cell line were further considered. The 40 top ranked genes were perturbed by siRNA in the well characterized human A431 squamous carcinoma cell line (Table S2). Of the selected genes, 31 are transcriptional regulators, 10 are oncogenes, and 7 are tumor suppressors.

RNA silencing experiments were carried out to knock down the expression of each individual gene, whereafter the expression of all genes in response to the perturbation was measured. The experiments were carried out with three biological replicates per gene. At steady-state, gene expression was measured using high-throughput RT-qPCR, totalling 18432 quantifications. Most targeted genes were seen to have dramatic reduction in expression, generally a stronger effect than for the other genes (Fig. S2). In fact 31 targets were significantly downregulated (p < 0.1 and log_2_ fold change < −2).

The perturbation response gene expression data were used for network inference with three methods, LASSO, LSCO, and TLSCO, each run in conjunction with nested bootstrapping. Each GRN inference method was run with varying parameters to produce GRNs in a range of different sparsities. Nested bootstrapping was then used to select the significantly supported links in each GRN, resulting in final sparsities that tend to never exceed 3-5 links/gene even if the natively inferred GRN had 40 links/gene, for example.

In order to select the GRN that has the links most likely to exist in reality, we compared how well each inferred GRN’s topology fits the data compared to shuffled topologies of the same GRN. The reason for validating the topology rather than the complete GRN including the actual parameter values is that those parameters are optimized to fit the data and therefore no suitable null model exists. Note that by topology we mean the structure of the GRN, i.e. the inferred links and their sign. For each topology, the parameters were fit to the observed data using crossvalidation, and the error was measured as the difference between predicted and observed values of the hold-out samples, after assembling the individual gene predictions into the full predicted matrix. This was done with the novel BalanceFitError (BFE) algorithm (Algorithm 1), which ensures that the error is balanced between sources, i.e. that the error is not merely pushed from the measurement to the process estimation or vice versa. Note that this algorithm is not a GRN inference method on its own but is rather a way to assess the quality of an inferred GRN.

Each inferred GRN was shuffled a hundred times, and the data was fit the same way to these topologies to estimate a null distribution of expected inference error. Note that since the parameters of the shuffled topologies are fit to minimize the error, this is a very stringent test of the inferred topology and for a suboptimal GRN one expects some of its shuffled topologies to by chance have lower error. Yet, several of the inferred GRNs greatly outperformed their null model, both when using the original training data (Fig. 2) and the independent validation data with double knockdown design in the same cell line (Fig. S3). We also calculated R^2^ values to show the proportion of the variation that our GRNs explain (Fig S6). All GRNs are available at https://dcolin.shinyapps.io/CancerGRN/. Five of the inferred topologies had 1000 times lower error than the median of the shuffled null model. The most accurate GRNs were inferred by Lasso, in terms of outperforming their null distributions. All but one GRN outperformed the median of their null distributions, and eight of the nineteen GRNs across all three methods outperformed all shuffled GRNs in their respective null distribution.

We also applied another null model to test how well the data fits the inferred GRN. Here we shuffled the original data one hundred times and fit these datasets to the inferred GRN in order to generate a null distribution. For many inferred GRNs the error was significantly lower than the median of the null distribution, both for the original training data (Fig. 3) and for independent validation data (Figs. 3 and S4).

The GRN that outperformed its null distribution by the largest margin was Bolasso_network_L1145_M115_support97.5_1.52e-03, which we will refer to as the best GRN, with 125 links, including 39 self-links, between 39 genes (Fig. 4) and a sparsity of 3.2 links/gene. The full name indicates certain properties, namely that 1145 links were natively inferred before NestBoot, 115 experiments were used, 97.5% bootstrap support was attained at FDR=0.05, and 1.52e-03 is the sparsity penalty parameter used. In this GRN’s overlap plot (Fig. 5) one can see the distributions of bootstrap values for both measured and shuffled data. The frequency of bootstrap support for measured data increases sharply at the right end above 98%, suggesting that this part of the distribution represents real and therefore highly reproducible links. In contrast, the shuffled data decreases towards support=1. The fact that some links inferred from shuffled data can attain such high bootstrap values can be attributed to the fact that the inference is done at a sparsity that yields very dense GRNs which may result in spurious links with high bootstrap support. However, the nested bootstrap framework monitors the distribution of spurious links and takes them into account when calculating FDR. The plot shows how FDR varies for different bootstrap support cutoffs.

**Fig. 4.**
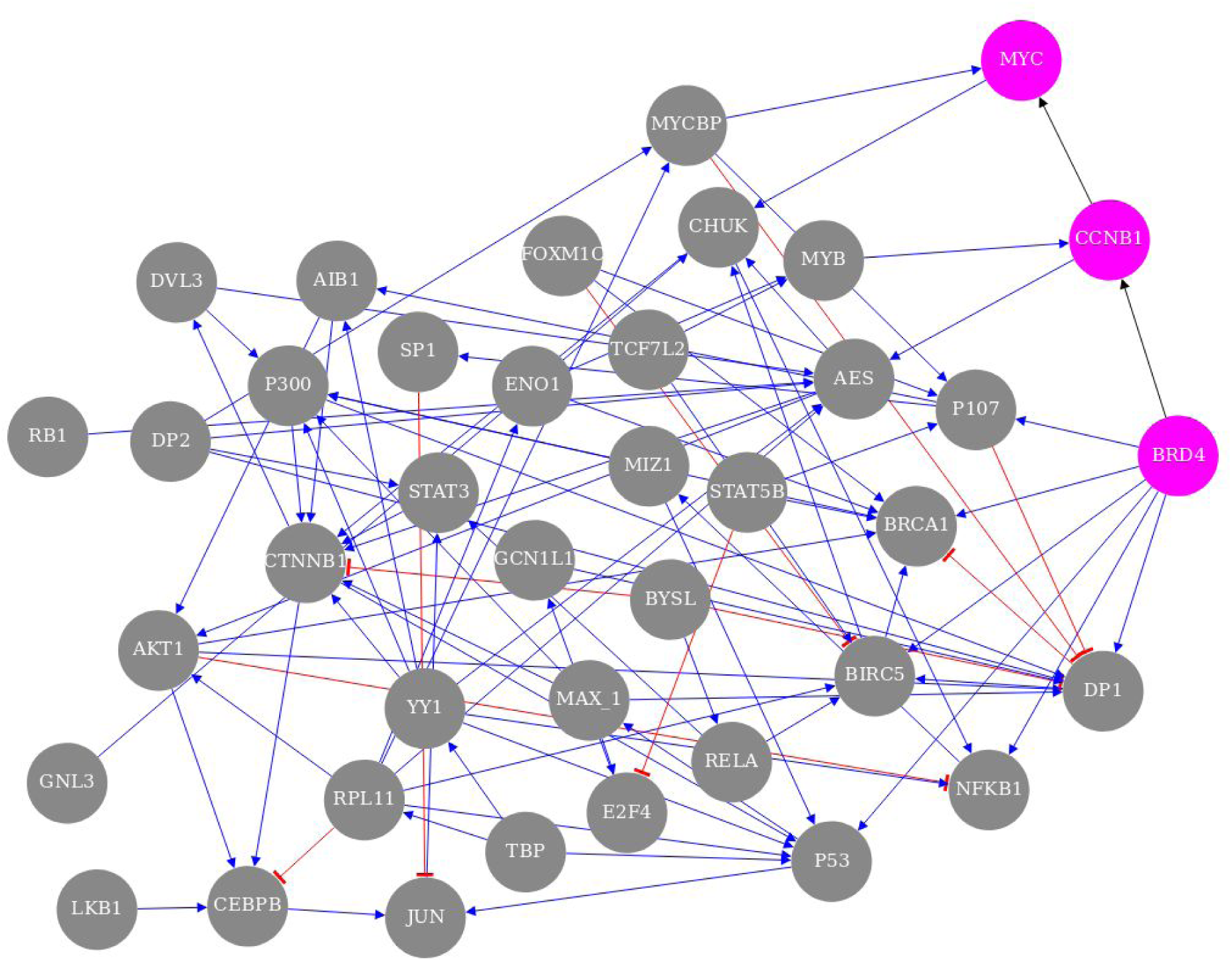
The overall best performing GRN. Each of the 125 links has at least 97.5% bootstrap support, and the sparsity is 3.2 links/gene among its 39 genes. The 39 self links are not shown. The genes involved in the external validation experiment, BDR4, CCNB1, and MYC, are highlighted pink. Blue links reflect up regulation while red reflect negative down regulation. The visualization was made by the provided shiny app.

**Fig. 5.**
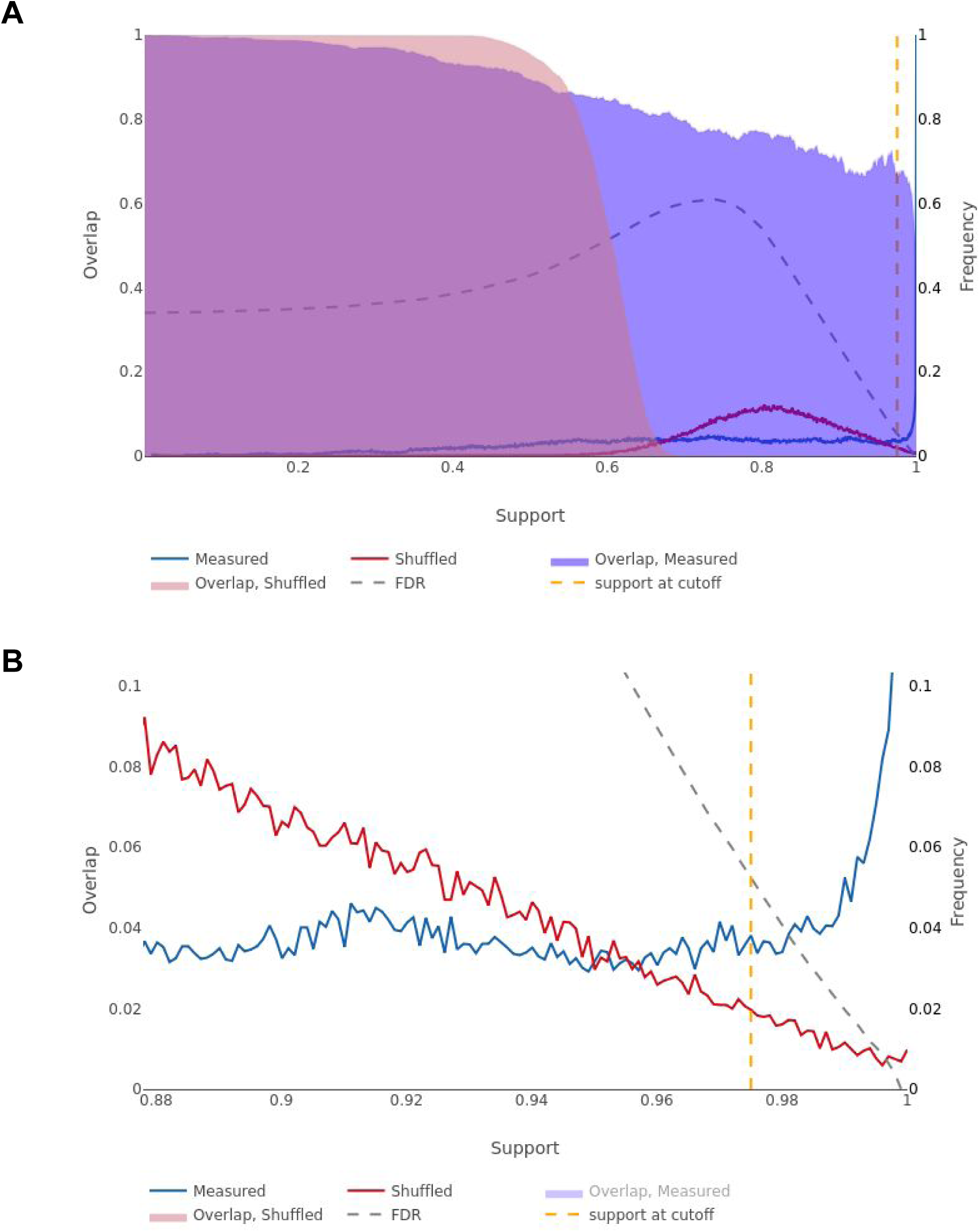
NestBoot output for the overall best performing GRN. **A** shows the entire bootstrap support range from 0 to 1, as well as overlap between all bootstrap GRNs for measured (blue) and shuffled (red) data. The FDR is estimated via a null background model based on networks inferred from shuffled data. This is done to restrict inclusion of false links by setting FDR e.g. to 5%. The dashed orange line represents the cutoff where this is reached, here at 97.5% bootstrap support. The dashed grey line shows how the FDR behaves as a function of the bootstrap support. **B** shows the fine detail of the curves in the support levels between 0.9 and 1. These visualizations were made by the provided shiny app.

One can also see the level of overlap between one hundred nested bootstrap runs in Fig. 5. Each run yields a bootstrap support for every link, which can be converted to a GRN for a given cutoff. For the measured data, the overlap (Jaccard) between runs stays relatively high (0.6) all the way to links with 100% bootstrap support, indicating that the reproducibility is high. In contrast, for the shuffled data not a single link with bootstrap support above 70% overlaps with another nested run, indicating poor reproducibility despite relatively high bootstrap support.

### Validation of the best GRN

Of the 125 links inferred in the top performing GRN, two novel MYC-related links were experimentally validated. The novel regulatory relationships BRD4→CCNB1 and CCNB1→MYC (Fig. 3) were examined in an independent study [28] in which JQ1 was used to inhibit BRD4 in the GTML2 cell line, a mouse brain tumor cell line that overexpresses human MYCN. The inferred activation of CCNB1 by BRD4 was supported by a significant reduction of CCNB1 expression when BRD4 was inhibited, from 7.12 to 7.04 average log(CPM) after 6h (Fig. S5). However, in order to study immediate effects of BRD4 inhibition we here performed a new analysis in which GTML2 cells were treated with JQ1 for just 2 hours. Again, CCNB1 expression was significantly reduced, from 7.48 to 7.04 average log(CPM). Support for the inferred activation of MYC by CCNB1 was found by the fact that the CCNB1 expression changed from normal newborn mouse brain to adult mouse brain (from FPKM 20.3 to 0.2) which agrees with the change of MYC (from FPKM 9.2 to 0.8) [28].

These validation experiments support a novel mechanism for MYC regulation inferred in the best GRN. While it is well known that BRD4 can activate MYC in some cancer types [29], the best GRN presents a regulatory route that goes via CCNB1 (Cyclin B1). Bound with cyclin-dependent protein kinases, CCNB1 is involved in controlling the cell cycle at mitosis. The findings here suggest that CCNB1’s role in regulating biological processes such as proliferation and oncogenesis can proceed via the activation of MYC.

Another type of validation is comparison to known links in public network resources. The links in the best GRN were searched for in the databases TRRUST [30], FunCoup [31], HumanNet [32], and STRING [33] as well as in our prior network from data mining. Where these reference networks contained undirected links, we compared them to an undirected version of our GRN. Many known interactions were witnessed in the best GRN (21 recovered from STRING), speaking to its ability to accurately infer what is known. The overlap with the other GRNs was significant (p < 0.1) in a hypergeometric test in all cases but one (Table S3).

## Discussion

This study carries out a complete workflow for inferring reliable GRNs, from the selection of genes, experimental perturbation, data collection, and GRN inference, to validation of the inferred GRNs. It was applied to 40 cancer-related genes whose GRN was studied in a human squamous carcinoma cell line. The collected dataset was used to infer variously sparse GRN using three inference techniques within the NestBoot framework. The predictiveness of the inferred GRNs was estimated using the novel BalanceFitError algorithm under crossvalidation. Almost all inferred GRNs were more predictive than expected by chance, and some were vastly more predictive. These top performing GRN were also able to predict an independent pairwise-gene perturbation validation dataset significantly better than expected by chance. The best GRN contains many known links as well as proposes many novel links, two of which were verified experimentally.

The performed gene perturbations caused a range of fold changes in both targeted and readout genes (Fig. S2). Knockdown is advantageous for GRN inference compared to complete inhibition through knockout, as that could alter the gene functioning within the cells to such an extent as to potentially drive the cell to any number of non-native states by activating alternative pathways to cope with the loss of the knocked out one. This would result in measurement of an altogether different cellular GRN which lacks the knocked out gene. With knockdown, a gene’s effect is lowered in the hope of measuring an otherwise wild-type GRN from the perspective of the single gene perturbation, across the gene repertoire.

The knockdown efficiency of each siRNA is unknown, and varies between genes. It may seem desirable to know the siRNA efficiency since this is a parameter in the perturbation design matrix that is used in the mathematical modelling. However, its value does not affect the inferred GRN’s topology, and since the topology is the main outcome of the inference, and what we compare to null, this lack of knowledge is inconsequential. Prior information, whether literature-curated, ChIP-seq, ATAC-seq has been shown to be of value in modern GRN investigations, and may also be helpful. Such integration could be built into the model as a method of constraining spurious link additions much the same as NestBoot restricts links based on shuffled link distributions. The NestBoot algorithm produces substantial accuracy improvement and we would anticipate further accuracy improvements from the addition of priors. However, such experimental information is not available for this study and we therefore pursued a strictly data driven approach.

Despite our efforts to measure absolute mRNA levels using spiked-in RNA as qPCR reference, MYC was not found to be a universal amplifier as previously claimed [34–36]. Our observation agrees with the results of [37]. In both their and our study, measurements were done after 72 hours. It is possible that MYC knockdown activates a response leading to rapid restoration of MYC expression, so that cells return to their original state within that time span, instead of reaching a new steady state. In our study, the targeted MYC transcript was not significantly repressed by the MYC siRNA. This may be caused by its unusually high turnover rate [38], which can make it difficult to knock down with siRNA [39]. The same lack of observed knockdown for the target was noted for three other transcription factors: SP1, LMYC, and JUN. This seems to suggest a need for optimizing the experimental protocol to obtain perturbed steady state conditions when knocking down certain transcription factors.

During the NestBoot procedure the sparsity of the native GRNs is varied from almost a full to almost an empty network. However, as NestBoot selects the strongest supported links only, the sparsity of the GRNs output by NestBoot does not vary much for the denser GRNs, and even less in gene makeup. We observe consistency across different sparsities, i.e. the smaller GRNs are mostly a subset of the larger ones. This consistency among sparsities adds further confidence beyond the GRNs’ predictiveness relative to a null distribution of shuffled topologies. Selecting the GRN with the optimal sparsity can be done in several ways. Here we followed the strategy of selecting the GRN with the best combination of coverage and predictiveness. Another criterion to select GRNs is the biological rationale that natural systems usually contain 3-5 links per gene[8,21].

In this study we face the problem of how to measure accuracy in the absence of a true network. Lacking such a gold standard it is impossible to determine if an inferred link is true or false. Instead, we compared each inferred network to a null distribution of GRNs with the same sparsity and indegree distribution. Since the prediction error depends on the weights of the links, it is crucial to fit each shuffled-link GRN to the data to give it reasonable weight estimates. To make the comparison fair, both the inferred GRN and the shuffled-link GRNs are refit to the data. By showing that the inferred GRN outperform their shuffled counterparts in terms of ability to explain the data, measured both by the wRSS and R^2^, we know that they have a topology closer to the unknown real GRN. The exact same procedure can then be applied to other data, such as the independent validation dataset. With enough repeated shuffled-link GRNs to produce a sufficient null distribution, this results in an unbiased estimate of how predictive a given GRN is compared to what is expected, despite lacking a known gold standard network. Benchmarking on data with a known gold standard shows that increased predictiveness measured this way generally agrees with higher accuracy.

## Acknowledgements

We thank S. Nelander and L.-G. Larsson for helpful discussions. The work of TEMN was in part supported by Swedish strategic research program eSSENCE, Sweden, 2-year post-doctoral fellowship, the National Cheng Kung University, Taiwan, and Ministry of Science and Technology of Taiwan [MOST 105-2218-E-006-016-MY2, 107-2634-F-006-009, 108-2634-F-006-009, 108-2218-E-006-046].

## References

1. Shulman LP. Analysis of microarray experiments of gene expression profiling. Yearbook of Obstetrics, Gynecology and Women’s Health. 2007;2007: 58–59.

2. Haury A-C, Mordelet F, Vera-Licona P, Vert J-P. TIGRESS: Trustful Inference of Gene REgulation using Stability Selection. BMC Syst Biol. 2012;6: 145.

3. Mordelet F, Vert J-P. SIRENE: supervised inference of regulatory networks. Bioinformatics. 2008;24: i76–82.

4. Guo S, Jiang Q, Chen L, Guo D. Gene regulatory network inference using PLS-based methods. BMC Bioinformatics. 2016;17: 545.

5. Tegnér J, Björkegren J. Perturbations to uncover gene networks. Trends in Genetics. 2007;23: 34–41.

6. Carro MS, Lim WK, Alvarez MJ, Bollo RJ, Zhao X, Snyder EY, et al. The transcriptional network for mesenchymal transformation of brain tumours. Nature. 2010;463: 318–325.

7. Olsen C, Fleming K, Prendergast N, Rubio R, Emmert-Streib F, Bontempi G, et al. Inference and validation of predictive gene networks from biomedical literature and gene expression data. Genomics. 2014;103: 329–336.

8. Marbach D, Costello JC, Küffner R, Vega NM, Prill RJ, Camacho DM, et al. Wisdom of crowds for robust gene network inference. Nat Methods. 2012;9: 796–804.

9. Castro DM, de Veaux NR, Miraldi ER, Bonneau R. Multi-study inference of regulatory networks for more accurate models of gene regulation. PLoS Comput Biol. 2019;15: e1006591.

10. Banf M, Zhao K, Rhee SY. METACLUSTER-an R package for context-specific expression analysis of metabolic gene clusters. Bioinformatics. 2019;35: 3178–3180.

11. Wani N, Raza K. Integrative approaches to reconstruct regulatory networks from multi-omics data: A review of state-of-the-art methods. Comput Biol Chem. 2019;83: 107120.

12. Delgado FM, Gómez-Vela F. Computational methods for Gene Regulatory Networks reconstruction and analysis: A review. Artificial Intelligence in Medicine. 2019. pp. 133–145. doi: 10.1016/j.artmed.2018.10.006

13. Bellot P, Olsen C, Salembier P, Oliveras-Vergés A, Meyer PE. NetBenchmark: a bioconductor package for reproducible benchmarks of gene regulatory network inference. BMC Bioinformatics. 2015;16: 312.

14. Schaffter T, Marbach D, Floreano D. GeneNetWeaver: in silico benchmark generation and performance profiling of network inference methods. Bioinformatics. 2011;27: 2263–2270.

15. Tjärnberg A, Morgan DC, Studham M, Nordling TEM, Sonnhammer ELL. GeneSPIDER - gene regulatory network inference benchmarking with controlled network and data properties. Mol Biosyst. 2017;13: 1304–1312.

16. Morgan D, Tjärnberg A, Nordling TEM, Sonnhammer ELL. A Generalized Framework for Controlling FDR in Gene Regulatory Network Inference. Bioinformatics. 2018. doi: 10.1093/bioinformatics/bty764

17. Gardner TS, di Bernardo D, Lorenz D, Collins JJ. Inferring genetic networks and identifying compound mode of action via expression profiling. Science. 2003;301: 102–105.

18. Bonneau R, Reiss DJ, Shannon P, Facciotti M, Hood L, Baliga NS, et al. The Inferelator: an algorithm for learning parsimonious regulatory networks from systems-biology data sets de novo. Genome Biol. 2006;7: R36.

19. Tjärnberg A, Nordling TEM, Studham M, Nelander S, Sonnhammer ELL. Avoiding pitfalls in L1-regularised inference of gene networks. Mol Biosyst. 2015;11: 287–296.

20. Gobbi A, Iorio F, Dawson KJ, Wedge DC, Tamborero D, Alexandrov LB, et al. Fast randomization of large genomic datasets while preserving alteration counts. Bioinformatics. 2014;30: i617–23.

21. Nordling T. Robust inference of gene regulatory networks. Jacobsen E, editor. PhD, KTH Royal Institute of Technology. 2013.

22. Tjärnberg A, Nordling TEM, Studham M, Sonnhammer ELL. Optimal sparsity criteria for network inference. J Comput Biol. 2013;20: 398–408.

23. Tibshirani R. Regression Shrinkage and Selection Via the Lasso. Journal of the Royal Statistical Society: Series B (Methodological). 1996. pp. 267–288. doi: 10.1111/j.2517-6161.1996.tb02080.x

24. de Groen PPN. An introduction to total least squares. 1998.

25. Michael Grant and Stephen Boyd. CVX: Matlab Software for Disciplined Convex Programming, version 2.1. mar, 2014. Available:\url{http://cvxr.com/cvx}

26. Cantone I, Marucci L, Iorio F, Ricci MA, Belcastro V, Bansal M, et al. A yeast synthetic network for in vivo assessment of reverse-engineering and modeling approaches. Cell. 2009;137: 172–181.

27. Chang LY, Pollard NS. Constrained least-squares optimization for robust estimation of center of rotation. J Biomech. 2007;40: 1392–1400.

28. Bolin S, Borgenvik A, Persson CU, Sundström A, Qi J, Bradner JE, et al. Combined BET bromodomain and CDK2 inhibition in MYC-driven medulloblastoma. Oncogene. 2018;37: 2850–2862.

29. Shi J, Wang Y, Zeng L, Wu Y, Deng J, Zhang Q, et al. Disrupting the interaction of BRD4 with diacetylated Twist suppresses tumorigenesis in basal-like breast cancer. Cancer Cell. 2014;25: 210–225.

30. Han H, Cho J-W, Lee S, Yun A, Kim H, Bae D, et al. TRRUST v2: an expanded reference database of human and mouse transcriptional regulatory interactions. Nucleic Acids Res. 2018;46: D380–D386.

31. Ogris C, Guala D, Sonnhammer ELL. FunCoup 4: new species, data, and visualization. Nucleic Acids Res. 2018;46: D601–D607.

32. Zeng X, Lin J, Lin C, Liu X, Rodriguez-Paton A. Structural Hole Spanner in HumanNet Identifies Disease Gene and Drug targets. IEEE Access. 2018; 1–1.

33. Szklarczyk D, Morris JH, Cook H, Kuhn M, Wyder S, Simonovic M, et al. The STRING database in 2017: quality-controlled protein-protein association networks, made broadly accessible. Nucleic Acids Res. 2017;45: D362–D368.

34. Lovén J, Orlando DA, Sigova AA, Lin CY, Rahl PB, Burge CB, et al. Revisiting global gene expression analysis. Cell. 2012;151: 476–482.

35. Lin CY, Lovén J, Rahl PB, Paranal RM, Burge CB, Bradner JE, et al. Transcriptional amplification in tumor cells with elevated c-Myc. Cell. 2012;151: 56–67.

36. Nie Z, Hu G, Wei G, Cui K, Yamane A, Resch W, et al. c-Myc is a universal amplifier of expressed genes in lymphocytes and embryonic stem cells. Cell. 2012;151: 68–79.

37. Nishiyama A, Sharov AA, Piao Y, Amano M, Amano T, Hoang HG, et al. Systematic repression of transcription factors reveals limited patterns of gene expression changes in ES cells. Sci Rep. 2013;3: 1390.

38. Jones TR, Cole MD. Rapid cytoplasmic turnover of c-myc mRNA: requirement of the 3’ untranslated sequences. Mol Cell Biol. 1987;7: 4513–4521.

39. Larsson E, Sander C, Marks D. mRNA turnover rate limits siRNA and microRNA efficacy. Mol Syst Biol. 2010;6: 433.

40. Hsieh AL, Walton ZE, Altman BJ, Stine ZE, Dang CV. MYC and metabolism on the path to cancer. Semin Cell Dev Biol. 2015;43: 11–21.

41. Ambion RNA-Seq Library Construction Kit. ThermoFisher; 01/2017. Available: http://tools.thermofisher.com/content/sfs/manuals/4452440C.pdf

42. CelluLyser TM Lysis and cDNA Synthesis Kit. TATTA; 10/2012. Available: http://www.tataa.com/wp-content/uploads/2012/10/prodblad_v03_tataa-CelluLyser.pdf

43. Biocenter T. TATAA Universal RNA Spike I. 2017 Oct. Available: https://webshop.tataa.com/dokument/Manual_TATAA%20Universal%20RNA%20Spike%20I%20SYBR-%20Probe_v1.3.pdf

44. Jitao David Zhang, Rudolf Biczok, and Markus Ruschhaupt. The ddCt Algorithm for the Analysis of Quantitative Real-Time PCR (qRT-PCR). 2017. Available: https://www.bioconductor.org/packages/release/bioc/html/ddCt.html

45. Delmore JE, Issa GC, Lemieux ME, Rahl PB, Shi J, Jacobs HM, et al. BET bromodomain inhibition as a therapeutic strategy to target c-Myc. Cell. 2011;146: 904–917.

46. Basso K, Margolin AA, Stolovitzky G, Klein U, Dalla-Favera R, Califano A. Reverse engineering of regulatory networks in human B cells. Nat Genet. 2005;37: 382–390.

47. Peck B, Ferber EC, Schulze A. Antagonism between FOXO and MYC Regulates Cellular Powerhouse. Front Oncol. 2013;3: 96.

48. Zwolinska AK, Heagle Whiting A, Beekman C, Sedivy JM, Marine J-C. Suppression of Myc oncogenic activity by nucleostemin haploinsufficiency. Oncogene. 2012;31: 3311–3321.

