## Supplemental Materials for "Perturbation-based gene regulatory network inference to unravel oncogenic mechanisms"

#### Supporting Information Legends

**Knockdown Data Collection**

**S1 Fig. Comparison of the two technical replicates in terms of qPCR cycles.**

**S2 Fig. Volcano plot describing fold change and significance of the knockdown responses.**

**S3 Fig. Independent validation of inferred GRNs' topologies.** Each x-axis tick mark shows the prediction performance in terms of the wRSS error of each inferred GRN topology (circles) fit to independent validation data under crossvalidation, compared to its shuffled topologies. The box displays the median and interquartile range, and whiskers bound points maximally extending 1.5 times this range. Beyond this, outlier points are shown.

**S4 Fig. Independent validation of inferred GRNs' fit to measured data.** Each x-axis tick mark shows the prediction performance in terms of the wRSS of an inferred GRN topology fit to independent validation data under crossvalidation, compared to its ability to fit shuffled data. X marks represent the inferred GRNs. The filled color box displays the median and interquartile range, and whiskers bound points maximally extending 1.5 times this range. Beyond this, outlier points are shown.

**S5 Fig. Experimental validation of the predicted activation of CCNB1 by BRD4.** CCNB1 expression was measured as $\log_{10}$ of counts per million reads after 2 and 6 hours of exposure to control (DMSO) and JQ1, which inhibits BRD4. The effect on CCNB1 was significant at both time points (p=0.048 and 0.014 by t-test).

**S6 Fig. Predictiveness of inferred GRN topologies using $R^2$ for the same GRNs as in Fig 2.**

### Supplemental

#### Knockdown Data Collection

A set of genes was assembled from different pathways and complexes, each interacting to some degree with the oncogene MYC [40] (Table S2). Each readout gene is perturbed in the human squamous carcinoma cell line A431 via transfection with short interfering RNAs (siRNAs). We then harvest, purify and prepare libraries using the Ambion Library Construction Kit[41]. A precise record of perturbations is key to modeling (next section). In order to minimize siRNA off target effects, two to three siRNA are used per target (Table S4), which are then averaged to purify the effects of the targeted siRNA perturbation. Cells were collected at 72 hours after siRNA knockdown and washed of Phosphate-Buffer Solution (PBS), and lysed using CelluLyser[42]. Cell counts were calculated using the resazurin fluorescence assay. Since no endogenous gene can be assumed to be free of MYC regulation, which is thought to be a universal transcriptional amplifier [34], a spike-in RNA transcript was added to each sample to act as a reference gene for the quantitative polymerase chain reaction (qPCR) analysis, added in proportion to the cell count before RNA isolation. It consisted of a 1000-base sequence with a 5' cap and a polyA tail. This was only used for normalization of mRNA level across samples[43]. Negative controls are included, which siRNA does not map to human genes, as well as an untreated control absent of any siRNA. The cDNA was prepared from the RNA and preamplified in preparation for the high-throughput qPCR screening. Finally, the transcript profiles with respect to the 40 genes were determined with TaqMan qPCR assays (Table S5) using Fluidigm Biomark 96x96 Dynamic Array integrated fluidic circuits. Raw qPCR output was then processed as a function of the samples' initial quantity returning a measure of fold change[44] via the experimental controls. Three experimental replicates were made per targeted perturbation and five outlying replicates were discarded due to clear machine read error, thus the dataset is composed of 40 genes (N) and 115 samples. Including all controls, a total of 18432 qPCRs were performed on 192 samples. Two technical replicates were performed to ensure minimal machine error. They generated very similar values up to 25 qPCR cycles (Fig. S1).

Experimental validation of individual interactions was performed on GTML2 brain tumor cells, which were cultured in serum-free stem cell medium as previously described [28] and treated for 2 hours with DMSO or JQ1 (500 nM). RNA was purified using the RNeasy Kit (Qiagen). RNA sequencing was performed using the Ion Proton™ System for Next-Generation Sequencing at NGI, SciLifeLab, Uppsala Biomedical Center (BMC), Sweden. All treatment conditions were performed in triplicates. All RNA sequence reads were processed and the differentially expressed genes were analyzed as previously described [28].

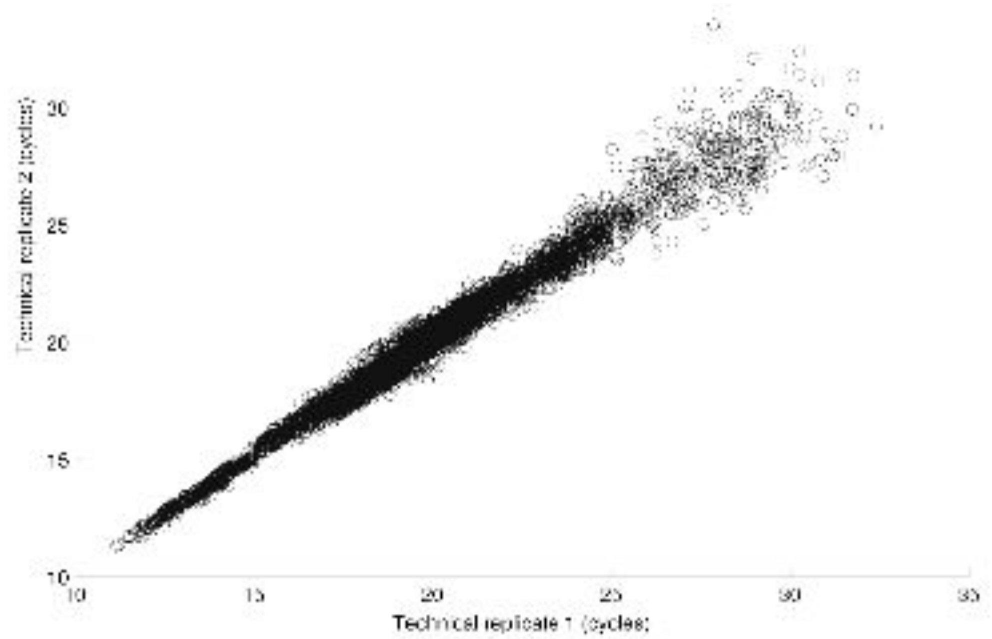

**S1 Fig.** Comparison of the two technical replicates in terms of qPCR cycles.

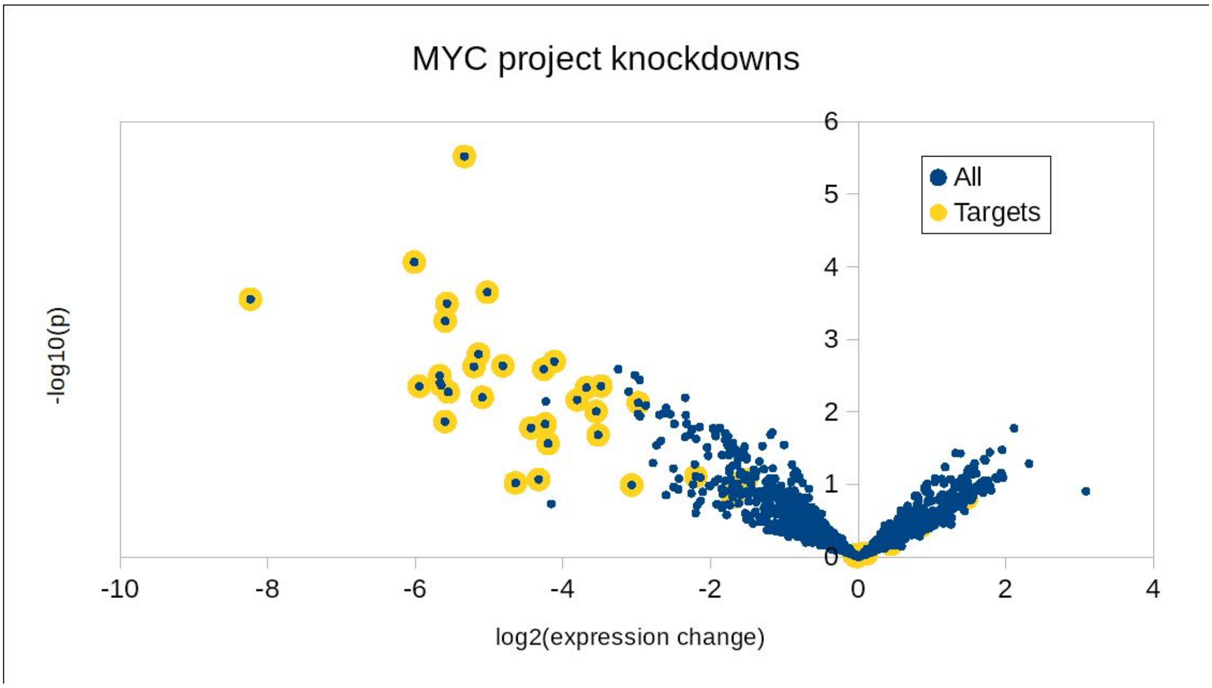

**S2 Fig.** Volcano plot describing fold change and significance of the knockdown responses.

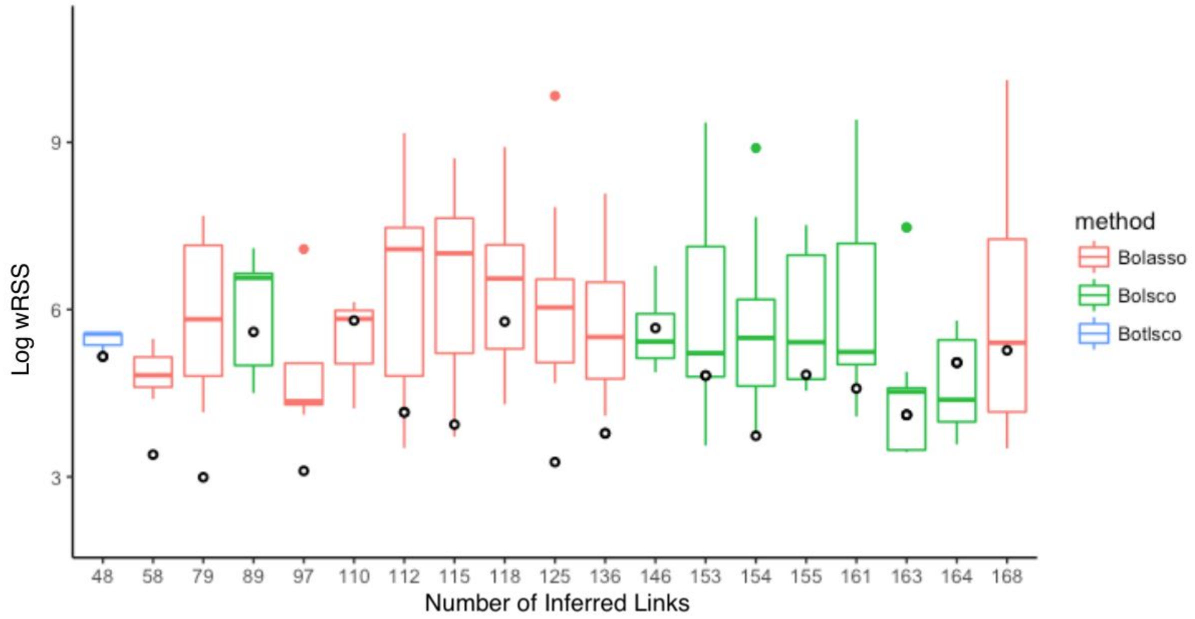

**S3 Fig.** Independent validation of inferred GRNs' topologies. Each x-axis tick mark shows the prediction performance in terms of the wRSS error of each inferred GRN topology (circles) fit to independent validation data under crossvalidation, compared to its shuffled topologies. The box displays the median and interquartile range, and whiskers bound points maximally extending 1.5 times this range. Beyond this, outlier points are shown.

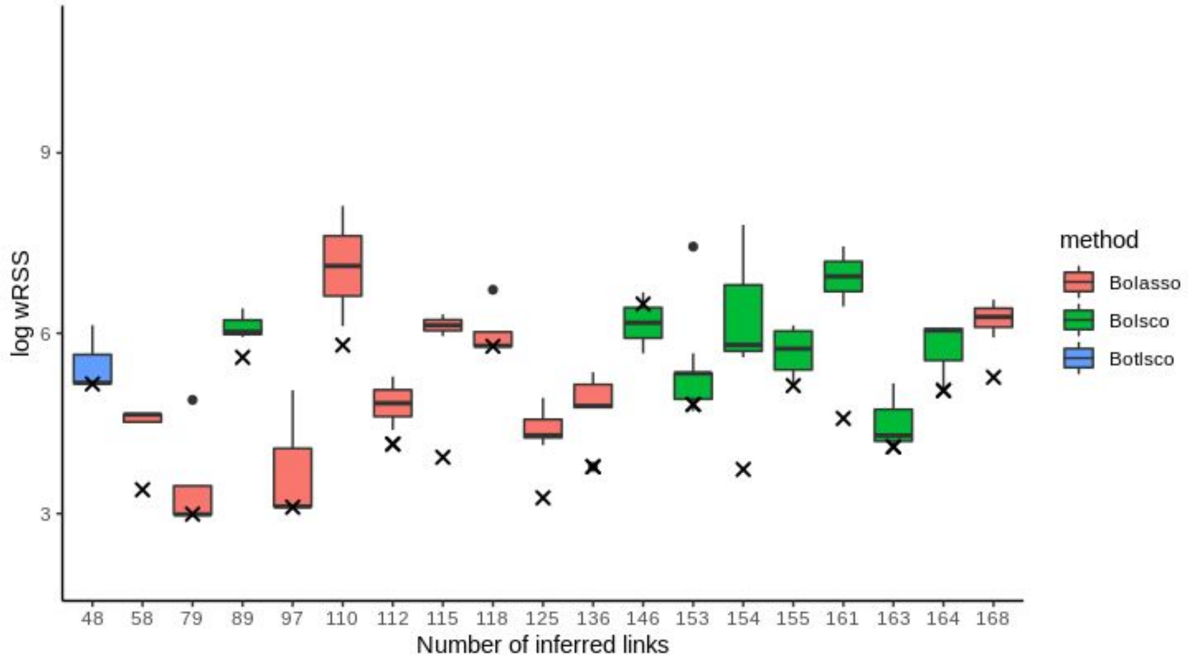

**S4 Fig.** Independent validation of inferred GRNs' fit to measured data. Each x-axis tick mark shows the prediction performance in terms of the wRSS of an inferred GRN topology fit to independent validation data under crossvalidation, compared to its ability to fit shuffled data. X marks represent the inferred GRNs. The filled color box displays the median and interquartile range, and whiskers bound points maximally extending 1.5 times this range. Beyond this, outlier points are shown.

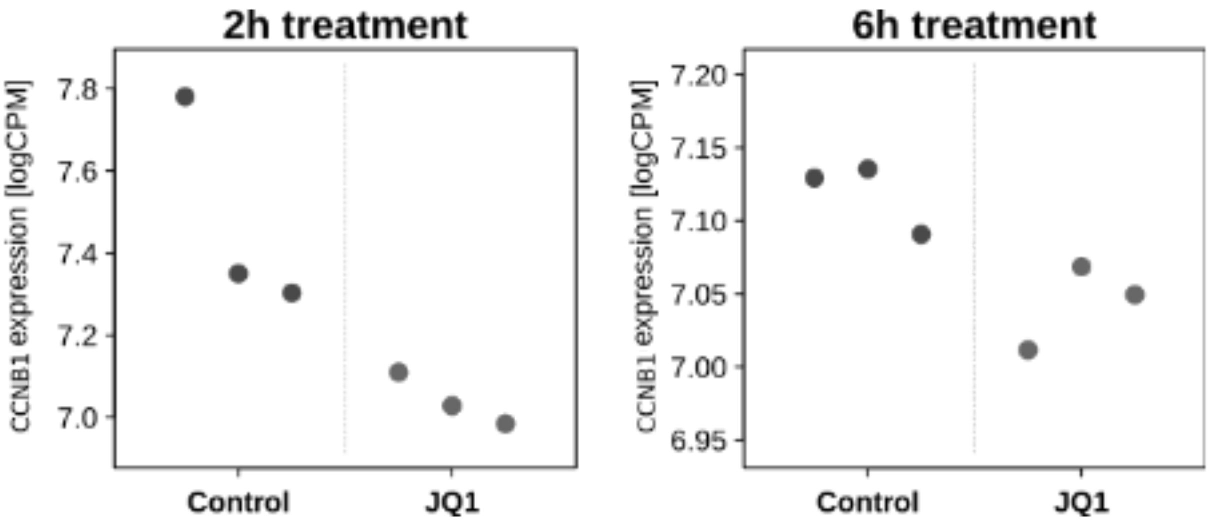

**S5 Fig.** Experimental validation of the predicted activation of CCNB1 by BRD4. CCNB1 expression was measured as $log_{10}$ of counts per million reads after 2 and 6 hours of exposure to control (DMSO) and JQ1, which inhibits BRD4. The effect on CCNB1 was significant at both time points (p=0.048 and 0.014 by t-test).

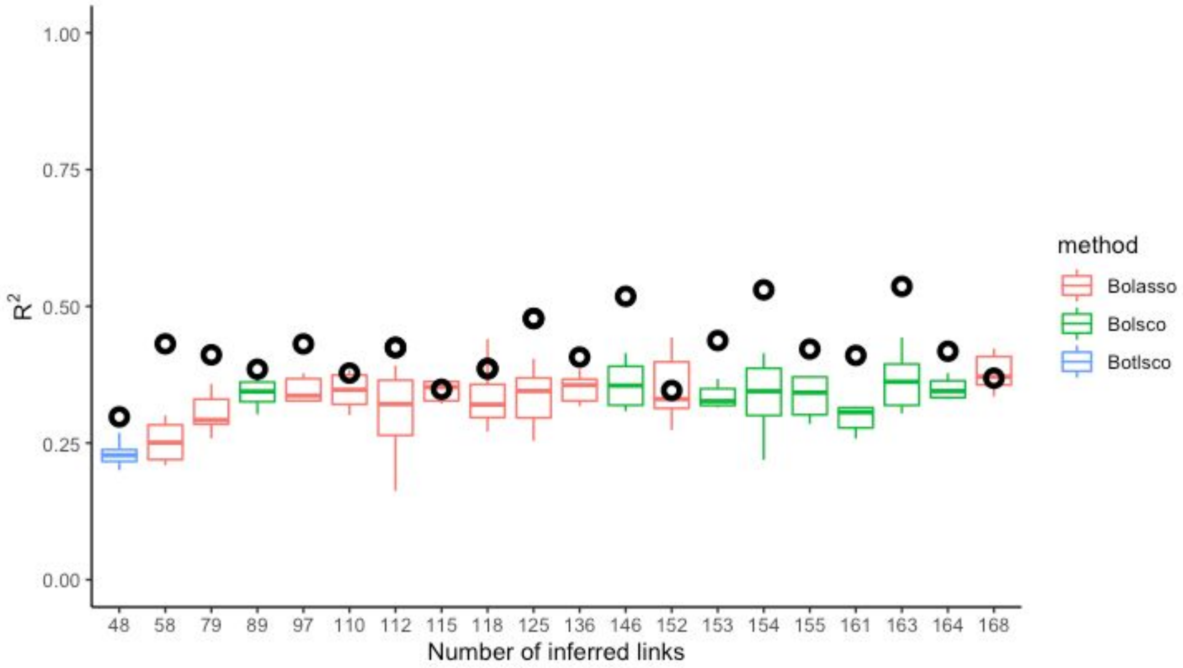

**S6 Fig.** Predictiveness of inferred GRN topologies using $R^2$ for the same GRNs as in Fig 2.

**S1 Table.** Description of matrix contents used in the linear model for GRN inference.

| **matrix** | **dimension (rows x columns)** | **contents** | **description/ value range** |
|---|---|---|---|
| Y | 40 x 115 | Gene expression | SNR=0.014, max=3.3, min=-8.7, condition number=55.3 |
| A | 40 x 40 | Interaction matrix (GRN) | Range dependent on BFE per init; initial inference returns variously sparse, weighted A, which is reweighted (see BFE section) |
| P | 40 x 115 | Perturbation/ design | Targeted perturbations (-1=KD, 0=no KD) |
| F | 40 x 115 | Process error | Range dependent on BFE per init |
| E | 40 x 115 | Measurement error | Range dependent on BFE per init |

**S2 Table**. Genes investigated. Description of perturbed genes in terms of pathways involved in, complexes formed, general notes (such as affect on Myc transcription), and RPKM expression in cell line A431. Pathway source material is the NCBI BioSystems Database unless otherwise noted. Complex information and General Notes are taken from NCBI. Gene names are listed as in our results and then synonyms in HGNC.

| Genes & synonyms | Pathway | Complex | Functional Annotation | RPKM |
|---|---|---|---|---|
| **AES** | ps1pathway | CtBP/CBP/TCF1/TLE1/AES, beta catenin/TCF1/CtBP/CBP/TLE1/AES/SMAD4 | Transcriptional corepressor/ regulation and signaling. First complex represses MYC, second activates MYC | 51.5 |
| **AIB1 / NCOA3** | Era genomic pathway | E2/ERA (dimer)/AIB1 | Transcriptional regulation and enzymatic behavior. Nuclear receptor coactivator. Complex activates MYC | 15.1 |
| **AKT1** | Il2 pi3kpathway | AKT1+plasma membrane | Kinase, enzyme, oncogene, metabolism. ser/thr kinase that activates MYC when in plasma membrane, oncogene | 13.2 |
| **BIRC5** | | | Transcription regulation. Protease inhibitor, Repressor | 44.1 |
| **BRCA1** | Myc repression pathway | | Transcription regulation,DNA-binding, metabolic and enzymatic activity, Tumor suppressor. Works w/MYC to repress psoriasin, tumor suppressor gene | 16.8 |
| **BRD4 / (M)CAP / HUNK(L)** | PI3K/ AKT pathway | activates Myc | Transcription regulation, metabolic and enzymatic activity, common complex w/P-TEFb (Delmore 2011[26]), oncogene | 8.1 |
| **BYSL** | | | [45,46] | 15.5 |
| **CCNB1** | FoxO signaling pathway | | metabolic activity, Cyclin B1<br>Ser/thr kinase activator | 145 |
| **CEBPB** | MEK/ERK signaling pathway | | Transcription factor, DNA-binding | 5.3 |

| | | | | |
|---|---|---|---|---|
| **CHUK** | | | Serine kinase, metabolic activity, | 14.6 |
| **CTNNB1** | BDNF signaling pathway<br>Wnt signaling pathway | | Transcription regulation, signaling, enzymatic activity,oncogene | 100.5 |
| **DP1** | e2f pathway | E2F1-3/DP | Transcription regulation, DNA-binding, complex activates MYC, oncogene, tumor suppressor gene | 6.6 |
| **DP2** | e2f pathway | E2F4/DP2/TRRAP, E2F4/DP2/GCN5 | Transcription regulation, DNA-binding, complexes activate MYC | 15.2 |
| **DVL3** | Beta catenin nuc pathway | TCF4/beta catenin/JUN/DVL3 | Signaling, complex activates Myc | 18.3 |
| **E2F4** | E2f pathway | E2F4/DP2/TRRAP, E2F4/DP2/GCN5, CEBPA/BRM/RB1/E2F4 | Transcription regulation, DNA-binding, First two complexes activate MYC, the second represses MYC | 15.2 |
| **ENO1** | Myc active pathway<br>Notch pathway | | Transcription regulation, DNA-binding, metabolic activity, represses MYC | 919.1 |
| **FOXO3A / FOXM1C** | Myc represions pathway | | Transcription regulation, DNA-binding, enzymatic activity, Tumor suppressor. [47] [47] | 13.3 |
| **GCN1(L1)** | | | [31] | 58.3 |
| **GNL3** | | | [48] [48] | 46.3 |
| **JUN*** | Ap1 pathway | JUN/JUND | activates as JUN/JUND | 15.3 |
| | Betacatenin nuc pathway | TCF4/beta catenin/JUN/DVL3 | Transcription regulation, DNA-binding, enzymatic activity, complex activates MYC, oncogene | |
| **LKB1 / STK11** | Lkb1 pathway | LKB1/ER alpha | complex activates MYC, Kinase, metabolic activity, Tumor suppressor. | 10 |
| **MAX** | Myc activation pathway | MYC/Max/RPL11,MYC/ Max/TRRAP/TIP60/TIP4 9A/TIP49B/BAF53 | | 21.7 |
| | Myc pathway | replication preinitiation complex, | Transcription regulation, DNA-binding, Tumor | |

| | | | | |
|---|---|---|---|---|
| | | MYC/Max/TRRAP/TIP60/TIP49A/TIP49B/BAF53 | suppressor. | |
| | Myc repression pathway | SMAD2-3/SMAD4/MYC/Max/MIZ-1,MYC/Max/NF-Y,MYC/Max/MIZ-1/ZO2, MYC/Max | | |
| **MIZ(-)1 / ZBTB17** | Myc pathway | MYC/Max/MIZ-1 | Transcription regulation, DNA-binding, Main MYC complex that represses | 2.5 |
| | Myc repression pathway | MIZ-1/p300,MYC/Max/MIZ-1,MYC/Max/MIZ-1/DNMT3A/GFI1 | | |
| **MYB** | Cmyb pathway | Myb, SKI/SIN3a/NCoR1/TIF1B/Myb | Transcription regulation, DNA-binding, metabolic and enzymatic activity, enzymatic activity, activates as Myb; represses MYC as part of complex, oncogene | 1.5 |
| **MYC** | Myc activation pathway | MYC/Max/RPL11,MYC/Max/TRRAP/TIP60/TIP49A/TIP49B/BAF53 | | 20.7 |
| | Myc pathway | MYC/Max/TRRAP/TIP60/TIP49A/TIP49B/BAF53, MYC/Max | Transcription regulation, DNA-binding, oncogene, | |
| | Myc repression pathway | MYC/Max/NF-Y,MYC/Max/MIZ-1/ZO2,MYC/Max | | |
| **MYCBP** | Notch pathway | MYCBP | Transcription regulation, binds to MYC, stimulates E-box transcription activation | 30.7 |
| **NFKB1** | Cd40 pathway | RelA/NFkappaB1 | Transcription regulation, DNA-binding, complex activates MYC | 15.9 |
| **P107 / PRB1 / RBL1** | Myc repression pathway | | works w/MYC/Max to repress MYC | 15.1 |
| **(E)P300** | Myc repression pathway | MIZ-1/p300 | Transcription regulation, metabolic activity, Tumor suppressor. Makes complex w/MIZ | 13.6 |
| **(T)P53** | Myc activation pathway | | Transcription regulation, DNA-binding, enzymatic activity, oncogene, tumor suppressor. | 26.9 |
| **RB1** | E2f pathway | CEBPA/BRM/RB1/E2F4 | Transcription regulation, DNA-binding, Tumor suppressor. complex represses MYC | 29.3 |
| **RELA** | Cd40 pathway | RelA/NFkappaB1 | Transcription regulation, | 19.1 |

| | | | | |
|---|---|---|---|---|
| | | | DNA-binding, complex activates MYC | |
| **RPL11** | Myc activation pathway | MYC/Max/RPL11 | | 716.1 |
| **SP1** | Foxm1 pathway | FOXM1C/SP1 | Transcription regulation, DNA-binding, complex works against MYC,tumor suppressor gene | 21.4 |
| | Myc repression pathway | SMAD2-3/SMAD4/SP1 | | |
| **STAT3** | Il6 7pathway | STAT3 dimer | Transcription regulation, DNA-binding, enzymatic activity, dimer activates MYC, oncogene | 19.9 |
| **STAT5 A/B** | Il2 stat5pathway | STAT5 dimer | Transcription regulation, DNA-binding, metabolic activity, dimer activates MYC (ENCODE[30] ), oncogene, tumor suppressor gene | 7.3 |
| **TBP** | Myc repression pathway | | Transcription regulation, DNA-binding, works against MYC in affecting multiple genes | 5.8 |
| **TCF7L2** | Wnt signaling pathway, beta catenin nuc pathway | TCF4/beta catenin/TNIK, TCF4/beta catenin/JUN/DVL3 | Transcription regulation, DNA-binding, enzymatic activity, signaling, complexes activate Myc (Sur 20[31] ), oncogene | 5.4 |
| **TIF1B / TRIM28** | Cmyb pathway | SKI/SIN3a/NCoR1/TIF1B /Myb | Transcription regulation, metabolic activity, part of complex that represses | 91 |
| **YY1** | Notch pathway | NICD/YY1 | Transcription regulation, DNA-binding, complex activates MYC | 27.9 |

**S3 Table.** Comparing the best GRN to reference networks. The networks include an expected prior network gathered from data mining, as well as the TRRUST v2, FunCoup v4, HumanNet v1 and STRING v10.5 databases. P-values are from a hypergeometric test. *links with confidence >= 0.90

| Network | Overlapping links | Links in DB | Links in GRN | p-value |
|---|---|---|---|---|
| Prior network | 5 | 40 | 86 | 0.08 |
| TRRUST | 7 | 132 | 86 | 0.66 |
| FunCoup | 14 | 146 | 76 | 0.01* |
| HumanNet | 9 | 81 | 76 | 0.02 |

| | | | | |
|---|---|---|---|---|
| STRING | 21 | 212 | 76 | 0.003 |

**S4 Table.** Used siRNA sequences

| Symbol | Entrez Gene | Ensembl Gene | Ambion siRNA ID | siRNA Sequences |
|---|---|---|---|---|
| AES | 166 | ENSG00000104964 | ASX053MJ | GCCUCAAGCUCGAAUGUGATT CAAAGACGAAUUUCAGCUATT GAACAUCGAGAUGCACAAATT |
| AIB1 | 8202 | ENSG00000124151 | ASX04YYH | CAGUAUAUCGAUUCUCGUUTT CAGUAAGACAGAUACGUCATT GGGCUUUUAUUGCGACCAATT |
| AKT1 | 207 | ENSG00000142208 | ASX04WP6 | GCGUGACCAUGAACGAGUUTT GAACAAUCCGAUUCACGUATT CGGUAGCACUUGACCUUUUTT |
| BIRC5 | 332 | ENSG00000089685 | ASX04ZH8 | GGACCACCGCAUCUCUACATT GCAGGUUCCUUAUCUGUCATT CAAAGGAAACCAACAAUAATT |
| BRCA1 | 672 | ENSG00000012048 | ASX053MG | GACCCAGUCUAUUAAAGAATT CAGCUACCCUUCCAUCAUATT CAUGCAACAUAACCUGAUATT |
| BRD4 | 23476 | ENSG00000141867 | ASX051I6 | UGAGCACAAUCAAGUCUAATT AGAUUGAAAUCGACUUUGATT CCUGAUUACUAUAAGAUCATT |
| BYSL | 705 | ENSG00000112578 | ASX055LN | AGGUGGUUGUGGACCCUGATT CCAGGAUUUUUGCCUCUAATT GGGAGGUAUUAUCUAAGUATT |
| CCNB1 | 891 | ENSG00000134057 | ASX04ZMX | CAACAUUACCUGUCAUAUATT GAAAUGUACCCUCCAGAAATT CACUUAUACUAAGCACCAATT |
| CEBPB | 1051 | ENSG00000172216 | ASX04ZP9 | CCGCCUGCCUUUAAAUCCATT GGCCCUGAGUAAUCGCUUATT GUAUAUUUUGGGAAUCUUUTT |
| CHUK | 1147 | ENSG00000213341 | ASX04WNH | GGACUAAAAGAAGACUAUATT GAAGGAUCCAAAGUGUAUATT GCCUAGAGCUAAGUACCAATT |
| CTNNB1 | 1499 | ENSG00000168036 | ASX04X6P | GGACCUAUACUUACGAAAATT GGAUGUUCACAACCGAAUUTT CUGUUGGAUUGAUUCGAAATT |
| DP1 | 7027 | ENSG00000198176 | ASX04XZ1 | CAUCUCCAAUGACAAAUUUTT GCAAGAAGACGGUCAUCGATT GCUCCAAUGGGUCUCAGUATT |
| DP2 | 7029 | ENSG00000114126 | ASX04XZ2 | GGUUAUGCUUGGAUGCAGATT GAUCCAUGAUGACAUAGAATT GGACUACUUCUGAACUCUATT |
| DVL3 | 1857 | ENSG00000161202 | ASX04X6U | GGUAAACGAGAUCAACUUUTT CCAGCUUCUUUGACUCAGATT CGGUCACUCUCAACAUGGATT |
| E2F4 | 1874 | ENSG00000205250 | ASX04ZWZ | AGAAAUCUUUGAUCCCACATT GUAUCGGGCUAAUCGAGAATT GGAUUUACGACAUUACCAATT |
| ENO1 | 2023 | ENSG00000074800 | ASX04ZYQ | CCGUGACCGAGUCUCUUCATT GCAGGUACAUCUCGCCUGATT CAGUGGUGUCUAUCGAAGATT |
| FOXM1C | 2305 | ENSG00000111206 | ASX0501W | GCUCAUACCUGGUACCUAUTT CACUAUCAACAAUAGCCUATT GGAUCAAGAUUAUUAACCATT |
| GCN1L1 | 10985 | ENSG00000089154 | ASX0518S | GGUGUAACCGAUUCACUAUTT GCAGCCUUGUUGCUCUUAATT GGCAGUUGAUUGUGGAUGATT |
| GNL3 | 26354 | ENSG00000163938 | ASX051NW | GGGUUGGAGUAAUUGGUUUTT GUAUUGUGGUAGACAUGAATT CCUCCGAUGUUGUCCUAGATT |
| JUN | 3725 | ENSG00000177606 | ASX050EK | GGCACAGCUUAAACAGAAATT GGAUCAAGGCGGAGAGGAATT CCAAGUGCCGAAAAAGGAATT |
| LKB1 | 6794 | ENSG00000118046 | ASX04WWS | GGCUCUUACGGCAAGGUGATT ACAUCACCACGGGUCUGUATT AGGAGGUUACGGCACAAAATT |
| MAX | 4149 | ENSG00000125952 | ASX050J3 | CAAACGGGCUCAUCAUAAUTT CACACACACCAGCAAGAUATT CAAAGACAGCUUUCACAGUTT |
| MYB | 4602 | ENSG00000118513 | ASX050LY | CCUCUCAUCUAGUAGAAGATT GGAAAAGCGAAUAAAGGAATT GAAUUGCUCCUAAUGUCAATT |
| MYC | 4609 | ENSG00000136997 | ASX050M5 | ACAGCCCACUGGUCCUCAATT GAGCUAAAACGGAGCUUUUTT AGACCUUCAUCAAAAACAUTT |
| MYCBP | 26292 | ENSG00000214114 | s25391 | GUAUGAAGCUAUUGUAGAATT UUCUACAAUAGCUUCAUACTT |
| NFKB1 | 4790 | ENSG00000109320 | ASX053S7 | GGCUCAUGUUUACAGCUUUTT CCACCUUCAUUCUCAACUUTT GCAGCUGUAUAAGUUACUATT |
| P107 | 5933 | ENSG00000080839 | ASX04XXC | GCUCUUUGCCUAUAGCUCATT GGAUGGACAUGUCAAAUCUTT CCACCAAAGUUACCACGAATT |
| P300 | 2033 | ENSG00000100393 | ASX04ZYU | GGACUACCCUAUCAAGUAATT CCACUACUGGAAUUCGGAATT GCCUGGUUAUAUAACCGGATT |
| P53 | 7157 | ENSG00000141510 | ASX053MH | GUAAUCUACUGGGACGGAATT GAAAUUUGCGUGUGGAGUATT GGUGAACCUUAGUACCUAATT |
| RB1 | 5925 | ENSG00000139687 | ASX04XXA | GGAUAGCAAAACAACUAGATT CCAGUACCAAAGUUGAUAATT GCGUGUAAAUUCUACUGCATT |
| RELA | 5970 | ENSG00000173039 | ASX04XXT | CCCUUUACGUCAUCCCUGATT GGAGUACCCUGAGGCUAUATT GGAUUGAGGAGAAACGUAATT |
| RPL11 | 6135 | ENSG00000142676 | ASX0571N | GGUGCGGGAGUAUGAGUUATT CAACUUCUCAGAUACUGGATT GGAACUUCGCAUCCGCAAATT |
| SP1 | 6667 | ENSG00000185591 | ASX04Y7O | GGCAGACCUUUACAACUCATT CCACAAGCCCAAACAAUCATT GCAACAUGGGAAUUAUGAATT |

| | | | | |
|---|---|---|---|---|
| STAT3 | 6774 | ENSG00000168610 | ASX04Y94 | GCACCUUCCUGCUAAGAUUTT GGCUGGACAAUAUCAUUGATT GCCUCAAGAUUGACCUAGATT |
| STAT5B | 6777 | ENSG00000173757 | ASX04Y97 | CACCCGCAAUGAUUACAGUTT GGAGACUUGAAUUACCUUATT GCAUCACCAUUGCUUGGAATT |
| TBP | 6908 | ENSG00000112592 | ASX04XTS | CCAACAAUUUAGUAGUUAUTT GCCUAUUCAGAACACCAAUTT CAGUGAAUCUUGGUUGUAATT |
| TCF7L2 | 6934 | ENSG00000148737 | ASX04XU9 | GAUGGAAGCUUACUAGAUUTT CAAUGAAUCAGAAACGAAUTT GGUCAACCAGUGUACCCAATT |
| TIF1B | 10155 | ENSG00000130726 | ASX05101 | GGCCCUAUUCUGUCACGAATT GCAACAUUGCAGAAGAGCATT GCGGAAAUGUGAGCGUGUATT |
| YY1 | 7528 | ENSG00000100811 | ASX053W2 | AGCUUUUGUUGAGAGUUCATT AGAACUCACCUCCUGAUUATT CAGAAUAUAUGACAGGAAATT |

**S5 Table.** TaqMan assays used for qPCR in Fluidigm Biomark

| TATAA Assay | Gene | TaqMan Assay |
|---|---|---|
| AES | AES | Hs01081011_g1 |
| AKT1 | AKT1 | Hs00920503_m1 |
| BCL2 | BCL2 | Hs00236808_s1 |
| BIRC5 | BIRC5 | Hs04194392_s1 |
| BRCA1 | BRCA1 | Hs01556193_m1 |
| BRD4 | BRD4 | Hs04188087_m1 |
| BYSL | BYSL | Hs00608266_m1 |
| CCNB1 | CCNB1 | Hs01030103_m1 |
| CEBPB | CEBPB | Hs00270923_s1 |
| CHUK | CHUK | Hs00989502_m1 |
| CTBP1 | CTBP | Hs00972284_m1 |
| CTNNB1 | CTNNB1 | Hs00170025_m1 |
| DVL3 | DVL3 | Hs00610263_m1 |
| E2F4 | E2F4 | Hs00608098_m1 |
| ENO1 | ENO1 | Hs00361415_m1 |
| EP300 | P300 | Hs00914223_m1 |
| FOXM1 | FOXM1C | Hs01073586_m1 |
| GCN1L1 | GCN1L1 | Hs00412445_m1 |
| GNL3 | GNL3 | Hs00205071_m1 |
| JUN | JUN | Hs01103582_s1 |
| JUND | JUND | Hs00534289_s1 |
| MAX_g1 | MAX | Hs00811069_g1 |
| MAX_m1 | MAX | Hs01105523_m1 |
| MYB | MYB | Hs00920556_m1 |
| MYC | MYC | Hs00153408_m1 |
| MYCBP-GJA+ | MYCBP | Hs00429315_g1 |
| MYCL | LMYC | Hs00607136_g1 |
| MYCL1 | LMYC | Hs00420495_m1 |
| MYCN | NMYC | Hs00232074_m1 |

| | | |
|---|---|---|
| NCOA3 | AIB1 | Hs00180722_m1 |
| NFKB1 | NFKB1 | Hs00765730_m1 |
| RB1 | RB1 | Hs01078066_m1 |
| RBL1 | P107 | Hs00765707_m1 |
| RELA | RELA | Hs01042010_m1 |
| RPL11 | RPL11 | Hs00831112_s1 |
| SP1 | SP1 | Hs00916521_m1 |
| STAT3 | STAT3 | Hs01047580_m1 |
| STAT5B | STAT5B | Hs00560035_m1 |
| STK11 | LKB1 | Hs00975988_m1 |
| TBP | TBP | Hs00427620_m1 |
| TCF7L2 | TCF7L2 | Hs01009044_m1 |
| TFDP1 | DP1 | Hs00955491_gH |
| TFDP2 | DP2 | Hs00963605_m1 |
| TP53 | P53 | Hs01034249_m1 |
| TRIM28 | TIF1B | Hs00232212_m1 |
| YY1 | YY1 | Hs00231533_m1 |
| ZBTB17 | MIZ1 | Hs01114794_g1 |
